# NAB2-STAT6 Fusion Proteins Drive Nuclear Condensate Formation and Transcriptional Reprogramming in Solitary Fibrous Tumors

**DOI:** 10.1101/2025.08.07.668994

**Authors:** Yi Li, Jose L. Mondaza-Hernandez, Gauthami Pulivendala, Fardous Elsenduny, Felipe Beckedorff, Guilherme M. Lavezzo, Farhan Vahdat Azad, Zikun Zhou, Jeremy Warren, Paulina I. Trevino, Clark A. Meyer, David Lombard, Javier Martin-Broto, David S. Moura, Heather N. Hayenga, Leonidas Bleris

## Abstract

Solitary fibrous tumor (SFT) is a rare and aggressive sarcoma driven by NAB2-STAT6 gene fusions, yet effective targeted therapies remain unavailable. Here, we report that the NAB2ex4-STAT6ex2 fusion variant forms nuclear condensates via liquid-liquid phase separation (LLPS) in engineered fibroblast models and primary SFT cells. These condensates co-localize with BRD4S and EGR1, key transcriptional regulators, and are functionally active, driving widespread transcriptional reprogramming. Treatment with Mithramycin A, a compound that disrupts EGR1-DNA interactions, dissolves NAB2-STAT6 condensates and reverses their aberrant gene expression and chromatin binding signatures. Our findings uncover a previously unrecognized role for NAB2-STAT6 in condensate-mediated oncogenic signaling and provide a mechanistic rationale for condensate-targeted therapy in SFT.

---

Solitary fibrous tumor (SFT) is a rare soft-tissue sarcoma (STS). The current long-term prognosis for patients with SFT is dismal, with a high rate of local recurrence and widespread metastases to the liver, bones, and lungs^1^. Currently, there are no FDA-approved systemic therapies that meaningfully improve long-term survival. Surgery or radiation is the first line of treatment against this cancer; however, this approach is location and time-limited^2^. In 2013, a defining discovery was made that all SFTs have a somatic intrachromosomal gene fusion between NAB2 and STAT6 on chromosome 12^3^. Although this single gene fusion appears to drive oncogenesis, therapeutic options and, most importantly, targeted studies and clinical trials remain limited. For unresectable disease, pazopanib is the most effective treatment in a phase II clinical trial^4^. It has a median progression-free survival of 5.6 months. Therefore, SFT remains an orphan malignancy in need of new mechanistic insights and effective therapeutic strategies.

It was recently reported that fusion oncoproteins can drive tumorigenesis by forming nuclear condensates via liquid-liquid phase separation (LLPS), which may result in reprogramming the *cis*-regulatory landscape by inducing *de novo* super-enhancers^5,6^. These condensates represent both a mechanistic insight and a therapeutic vulnerability, as their dissolution can reverse oncogenic transcriptional programs^7^. Herein, we hypothesized that NAB2-STAT6 fusion proteins may form nuclear condensates, and that targeting these condensates could lead to a new treatment avenue for SFT.

First, we observed that NAB2-STAT6 fusion proteins contain physicochemical features commonly associated with phase separation (intrinsically disordered regions/IDRs in NAB2ex4-STAT6ex2 fusion, **Figure 1a**). Indeed, using the computational fusion oncoprotein (FO)-Puncta ML model^8^, NAB2ex4-STAT6ex2 fusions were predicted to form condensates (Group 3: enrichment in histidine, serine, and proline amino acids, **Figure 1a**). To confirm the formation of NAB2-STAT6 fusion protein condensates in SFTs, we first prepared lentiviral vectors expressing the NAB2ex4-STAT6ex2 fusion construct conjugated with the red fluorescent protein, mKate. A truncated mKate-conjugated construct (NAB2ex2) was created for use as a negative control (only exons 1 and 2 of NAB2, **Figure S1**). The constructs of fusion proteins tagged with mKate were stably integrated into immortalized normal human lung fibroblasts, which were chosen for their similarity to the suspected origin of SFT, namely, mesenchymal stem cells of fibroblastic origin^1^. As shown in **Figure S1**, the control mKate-NAB2ex2 fusion proteins were mainly localized in the cytosol (**Lf-C2**). In contrast, confocal fluorescence microscopy analysis showed that NAB2ex4-STAT6ex2 fusion proteins formed round condensates (puncta) in the nucleus (**Figure 1b**, named **Lf-4-2**). This report is the first observation of NAB2-STAT6 fusion protein condensates in SFT cell models.

**Figure 1:**
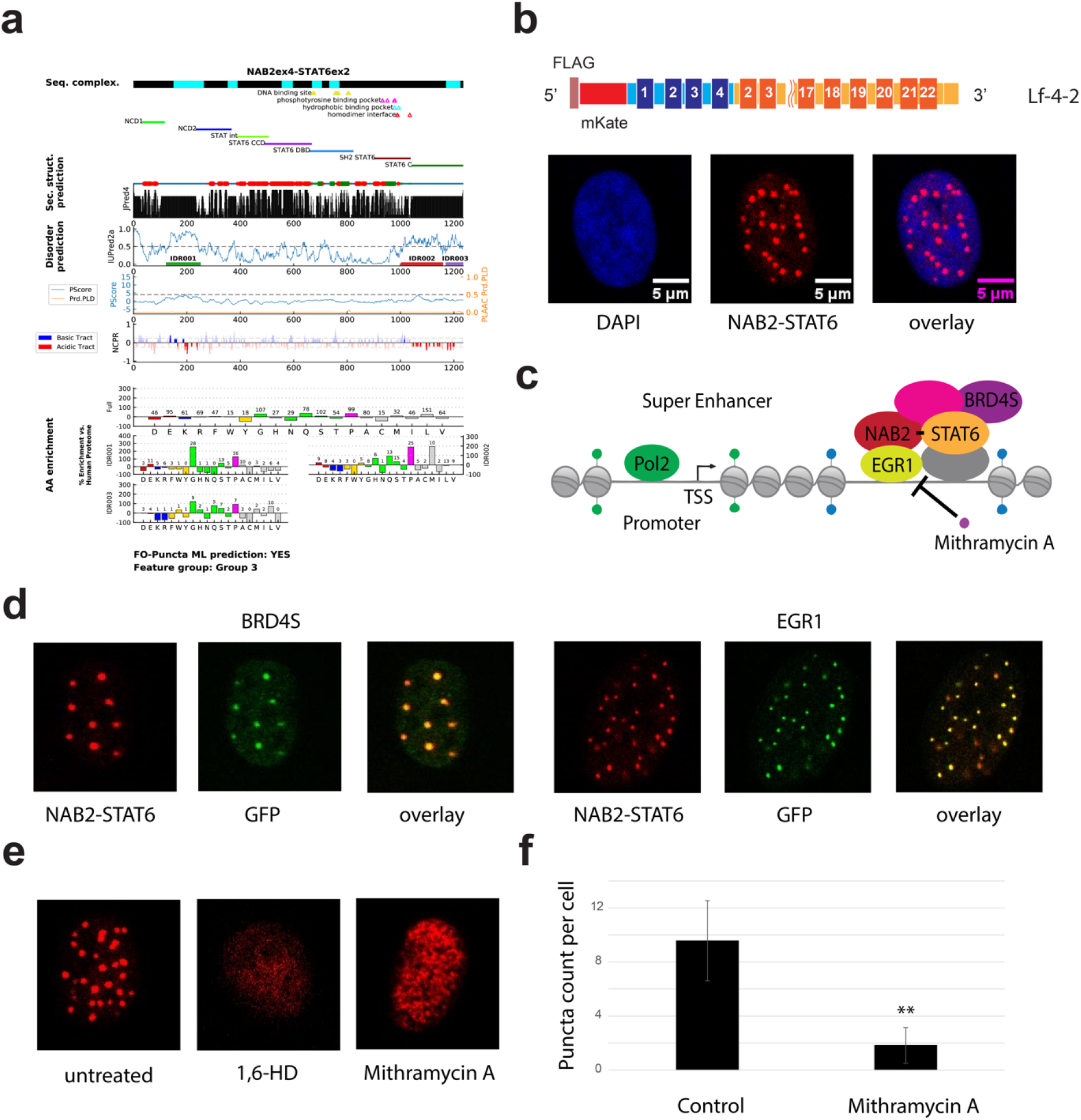
Mithramycin A dissolved NAB2ex4-STAT6ex2 condensates in modified immortalized human lung fibroblasts. **(a)** Prediction of NAB2ex4-STAT6ex2 condensate formation using the FO-Puncta ML model. **(b)** Confocal fluorescence microscopy (40X magnification) showed that NAB2ex4-STAT6ex2 fusion formed condensates in the nucleus. (red) mKate, (cyan) DAPI **(c)** Illustration of the proposed mechanism for forming NAB2-STAT6 condensates. **(d)** Confocal fluorescence microscopy (40X magnification) showed co-localization of NAB2ex4-STAT6ex2 (mKate) and BRD4S or EGR1 (GFP) in the nucleus. **(e)** 1.5% 1,6-hexanediol (10 minutes) or 300 nM Mithramycin A (24 hours) effectively dissolved NAB2ex4-STAT6ex2 condensates. **(f)** Treatment of 300 nM Mithramycin A reduced NAB2ex4-STAT6ex2 puncta count per cell by 81% (** indicated p-value < 0.01 using two-tailed, paired *t*-test).

Recent studies have identified bromodomain and extraterminal domain (BET) proteins as nucleation seeds for condensates^9^. Notably, the short isoform of bromodomain containing protein 4 BRD4 (BRD4S, which lacks the C-terminal extension of BRD4), has been shown to promote phase separation and the formation of nuclear puncta (**Figure 1c**)^10^. In parallel, NAB2ex4-STAT6ex2 fusions have been shown to exert transcriptional activation via interaction with EGR1 (early growth response 1)^11^. To investigate whether BRD4S or EGR1 co-localizes with NAB2-STAT6 condensates, we stably integrated GFP-BRD4S or EGR1-GFP into Lf-4-2 cells. As shown in **Figure 1d**, microscopy revealed extensive nuclear co-localization between BRD4S/EGR1 (green) and NAB2-STAT6 puncta (red). These results support a model in which BRD4S and EGR1 may act as functional components or cofactors within NAB2-STAT6-driven condensates, potentially controlling or amplifying their transcriptional regulatory effects.

Subsequently, given the established interaction between NAB2-STAT6 and EGR1, and the observed co-localization of these proteins within nuclear condensates, we hypothesized that disrupting EGR1’s binding to its GC-rich DNA target sites^12^, could interfere with the NAB2-STAT6 condensates. Mithramycin A, a clinically studied DNA-binding antibiotic known to block EGR1-DNA interactions, was selected as a candidate small-molecule disruptor.

Remarkably, treatment with Mithramycin A (300 nM) led to a dramatic reduction in the number of nuclear puncta formed by NAB2-STAT6 fusion proteins, comparable to the effect of 1,6-hexanediol^13^, a generic condensate-dissolving compound (**Figure 1e-f, Figure S2**, and **Table S1**, average numbers of puncta per cell: 9.6±3.0 in DMSO-treated cells and 1.8±1.3 in Mithramycin A-treated cells). Importantly, Mithramycin A-mediated NAB2-STAT6 condensate dissolution was not a byproduct of cell apoptosis, as minimal apoptosis was observed within 24 hours of Mithramycin A treatment (**Figure S3**). These results suggest that EGR1-DNA binding may be essential for the maintenance of NAB2-STAT6 condensates.

To determine whether Mithramycin A-mediated condensate dissolution can reverse aberrant NAB2-STAT6-induced gene expression, we performed bulk RNA sequencing in Lf-C2 and Lf-4-2 cells, each with or without treatment. As shown in **Figure 2a** (left: Venn diagram, middle: cells without the treatment) and **Table S2**, 1,154 genes were significantly upregulated in Lf-4-2 cells compared to Lf-C2 without Mithramycin A treatment (candidate NAB2ex4-STAT6ex2-inducing gene targets, **Figure S4a**). Subsequently, the protein class analysis using the PANTHER classification system revealed the enrichment of several protein classes (**Figure S4b**, metabolite interconversion enzymes: 148 genes, gene-specific transcriptional regulators: 76 genes)^14^.

**Figure 2:**
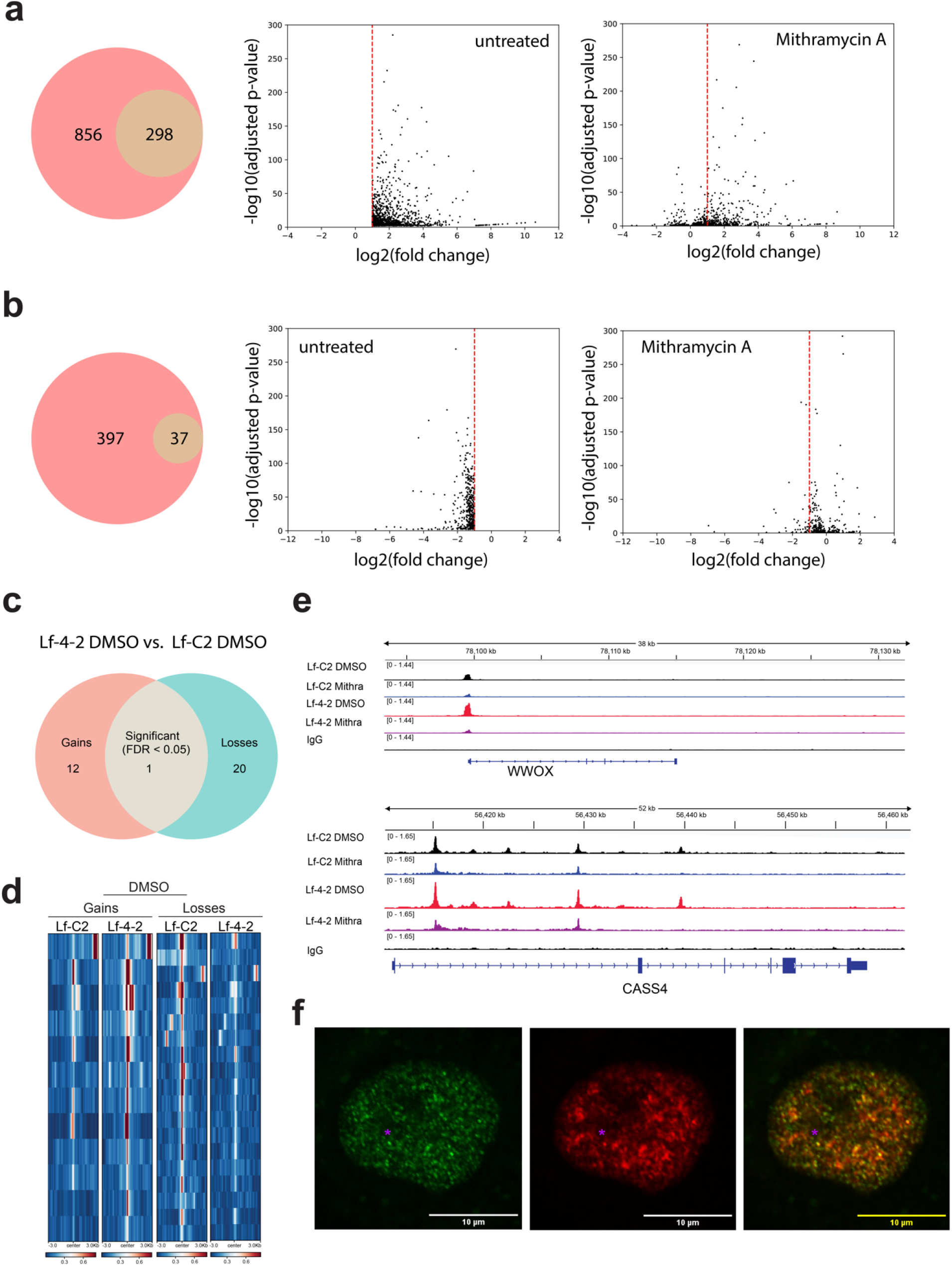
Mithramycin A reversed NAB2ex4-STAT6ex2-induced transcriptional signature and EGR1-binding patterns in immortalized human lung fibroblasts. **(a)** Treatment with Mithramycin A abolished the upregulation of 856 (74.2%) genes that were upregulated in Lf-4-2 cells compared to Lf-C2 cells. **(left)** Venn diagram, **(middle)** Volcano plot of genes upregulated in Lf-4-2 cells without Mithramycin A treatment, **(right)** Volcano plot of the same candidate genes upon Mithramycin A treatment. **(b)** Treatment with Mithramycin A abolished the downregulation of expression of 397 (91.5%) genes that were downregulated in Lf-4-2 cells compared to Lf-C2 cells. **(left)** Venn diagram, **(middle)** Volcano plot of genes downregulated in Lf-4-2 cells without Mithramycin A treatment, **(right)** Volcano plot of the same candidate genes upon Mithramycin A treatment. **(c)** Venn diagrams of shared/overlapping and unique peaks gained/lost as measured by ChIP-Seq for DMSO-treated Lf-4-2 vs. DMSO-treated Lf-C2. **(d)** Heatmap depicting genome-wide distribution of EGR1 peaks in DMSO-treated Lf-4-2 and DMSO-treated Lf-C2. **(e)** Integrative Genomics Viewer (IGV) tracks for CASS4 and WWOX locus in Lf-C2 and Lf-4-2 cells treated with either DMSO or Mithramycin A. **(f)** Immunofluorescence assay showed that NAB2/NAB2-STAT6 puncta (middle panel) overlapped with BRD4 puncta (left and right panels) in the primary NAB2ex4-STAT6ex2 SFT-739 cells (63X magnification).

Encouragingly, with RNA-seq using the same filtering criteria, Mithramycin A abolished the upregulation of expression of 856 (74.2%) candidate NAB2-STAT6-induced genes (**Figure 2a)**, indicating a broad reversal of the fusion-driven transcriptional program. For example, the expression of Interferon Induced Protein with Tetratricopeptide Repeats 1 (IFIT1)^15^, a cytoplasmic protein involved in Interferon-mediated antiviral responses, was upregulated in Lf-4-2 cells (**Table S3** and **Figure S5a**). In contrast, no significant overexpression of IFIT1 was observed in Mithramycin A-treated Lf-4-2 cells (**Table S3** and **Figure S5b**). Similar results were observed for another Interferon-mediated protein, IFIT2 (Interferon Induced Protein with Tetratricopeptide Repeats 2, **Figures S5c** and **S5d**)^15^. Additionally, among the 434 genes significantly downregulated in Lf-4-2 cells compared to Lf-C2 (**Table S4**), treatment with Mithramycin A abolished the downregulation of expression of 397 genes (91.5%, **Figure 2b**). These data demonstrate that NAB2-STAT6-induced condensates drive widespread, aberrant gene expression and that Mithramycin A disrupts this program at scale, restoring gene expression profiles toward those of non-fusion-expressing cells.

To directly assess whether Mithramycin A treatment disrupts NAB2-STAT6-dependent, EGR1-mediated chromatin binding, we performed chromatin immunoprecipitation followed by sequencing (ChIP-seq) in Lf-C2 and Lf-4-2 cells using an anti-human EGR1 antibody. As shown in **Figures 2c** and **2d**, compared to the control, NAB2ex4-STAT6ex2 expression induced 12 EGR1-target peaks (**Table S5**) and eliminated 20 peaks (**Table S6**). Importantly, upon treatment with Mithramycin A, 10 of the 12 gained peaks were lost (83.3%, **Table S5**, lost peaks highlighted in blue), supporting the hypothesis that Mithramycin A reverses EGR1-mediated, NAB2-STAT6-dependent binding at these loci (**Figure 1c**). Among the four protein-coding genes (Transcription Factor 25 (TCF25), Cas Scaffold Protein Family Member 4 (CASS4), WW Domain Containing Oxidoreductase (WWOX), and Cytidine 5’-Triphosphate Synthetase (CTPS1)) from the 10 identified peaks, two of them (CASS4 and WWOX, **Figure 2e** and **Figure S5**) were also identified by RNA-seq analysis as both NAB2-STAT6-dependent and Mithramycin A-sensitive gene targets (**Table S3**).

To validate our findings in a clinically relevant model, we next examined primary NAB2ex4-STAT6ex2-expressing SFT cells (SFT-739) using immunofluorescence. Antibodies targeting BRD4 and the N-terminal region of NAB2 were used to assess the presence and localization of nuclear condensates. As shown in **Figure 2f**, distinct NAB2/NAB2-STAT6 puncta features were detected in the primary SFT-739 cells (middle panel, asterisk), which, significantly, overlapped with the observed BRD4 puncta (right panel, asterisk). We emphasized that the more diffuse nuclear expression pattern for the anti-NAB2/NAB2-STAT6 antibody assay (**Figure 2f**, middle panel) may be due to wild-type NAB2 proteins, which are also recognized by this antibody and known to be expressed in the nucleus^16^. Nonetheless, the punctate co-localization of NAB2/NAB2-STAT6 and BRD4 in patient-derived cells confirms the physiological relevance of our findings.

It is worth noting that, depending on specific breakpoints within NAB2 and STAT6 genes, at least 6 different NAB2-STAT6 fusion types have been reported. Based on their clinicopathological features (e.g., mitotic figures, patient ages), these fusion types can be broadly classified into two groups, represented by NAB2ex4-STAT6ex2 and NAB2ex6-STAT6ex17^17^. Therefore, future studies will explore whether other NAB2-STAT6 fusion types also form nuclear condensates.

Beyond BRD4S and EGR1, we also plan to systemically explore the components (e.g., transcriptional and epigenetic regulators^18^) of NAB2-STAT6 condensates and their functions. For this, we noted that for NAB2ex4-STAT6ex2 condensates, Mithramycin A-induced dissolution preferentially abolished the upregulation of gene-specific transcriptional regulators (**Table S7**). It should be noted that mithramycin A is known to bind G-C-rich sequences in DNA, thereby inhibiting the interaction of specific transcription factors with the DNA, including EGR1^19,20^. Thus, some of the observed effects are likely mediated via direct inhibition of EGR1 activity.

Although Mithramycin A was historically withdrawn due to toxicity, several next-generation analogs, such as EC-8042 (AIT-102), have demonstrated comparable efficacy with significantly improved safety profiles^21^. These compounds merit further investigation in preclinical SFT models—either as monotherapy or in combination with standard-of-care drugs or compounds shown to be active in SFT, such as BET inhibitors^22^. In conclusion, our findings reveal that NAB2-STAT6 fusion proteins drive the formation of nuclear condensates in SFT, and that these structures are functionally active, pharmacologically targetable, and clinically relevant. This work deepens our understanding of the molecular underpinnings of SFT and lays the foundation for a novel condensate-based therapeutic strategy, with potential implications for other fusion-driven malignancies.

## Acknowledgements

We thank the laboratory members in the Li, Bleris, Hayenga, Lombard, and Martin-Broto labs for their support and discussions.

LB, JMB, HNH, CAM, DSM, and YL acknowledge funding from the US National Institutes of Health (NIH) grant 1R01CA283330. LB acknowledges funding from the Cecil H. and Ida Green Endowment at the University of Texas at Dallas. HNH acknowledges funding from the University of Texas at Dallas Bioengineering Transform Grant and the Vice President Accelerator Award. DSM is a recipient of a Miguel Servet contract financed by the National Institute of Health Carlos III (ISCIII) (CP24/00131). DBL’s laboratory is supported by R01CA253986 and R33AG077856, as well as the DoD (ME200030), Florida Department of Health (24B12), Melanoma Research Alliance (1434401), the Horowitz Solitary Fibrous Tumor initiative, and the Miami VA Healthcare System GRECC. For DBL, research reported in this publication was performed in part at the Onco-Genomics Shared Resource (OGSR) of the Sylvester Comprehensive Cancer Center at the University of Miami, RRID: SCR_022502, which is supported by the National Cancer Institute (NCI) of the National Institutes of Health (NIH) under award number P30CA240139. The content is solely the responsibility of the authors and does not necessarily represent the official views of the NIH.

## Data Availability

Data collected for this article are available in the **Supplemental Materials**.

## Materials and methods

### Mammalian cell culture

The hTERT-immortalized human lung fibroblast cell line (Lf) was obtained from the American Type Culture Collection (catalog #: CRL-4058) and maintained at 37°C, 100% humidity, and 5% CO_2_. Lf-C2 and Lf-4-2 cells were grown in Fibroblast Basal Medium (catalog #: PCS-201-030) supplemented with Fibroblast Growth Kit-Low serum (ATCC, catalog #: PCS-201-041) and 0.3 µg/mL of puromycin (Gibco, catalog #: A1113803). AAV-293 cells were maintained at 37°C, 100% humidity, and 5% CO_2_. The cells were grown in Dulbecco’s modified Eagle’s medium (Invitrogen, catalog #: 11965–1181) supplemented with 10% fetal bovine serum (Invitrogen, catalog #: 26140), 0.1 mM MEM non-essential amino acids (Invitrogen, catalog #: 11140–050), and 100 units/mL of penicillin and 100 µg/mL of streptomycin (Invitrogen, catalog #: 15140). To pass the cells, the adherent culture was first washed with PBS (Mediatech, catalog #: 21-030-CM), then trypsinized with Trypsin-EDTA (Invitrogen, catalog number: 25200) and finally diluted in fresh medium.

The SFT-739 cells were kindly provided by Dr. Florian Haller’s group (Comprehensive Cancer Center, Erlangen, Germany). Cells were maintained at 37°C, 100% humidity, and 5% CO2 in DMEM medium (Gibco, catalog #: 61965026) supplemented with 10% FBS (Gibco, catalog #: A5256701) and 100 units/mL of penicillin and 100 µg/mL of streptomycin (Gibco, catalog #: 15140122). Cells were passed once per week upon reaching confluency.

### Recombinant lentiviral vector production

AAV-293 cells were seeded at 70–80% confluency for lentiviral vector production. The cells were then transfected with NAB2ex4-STAT6ex2, VSV-G, and psPAX2 expression plasmids using JetPRIME (Polyplus Transfection, catalog #: 101000046). After overnight incubation, the media were replaced with complete DMEM growth media. Viral vectors were harvested 48-, 72-, and 96-hours post-transfection. The combined harvest was centrifuged at 1,250 rpm for 5 minutes, and the supernatant was transferred to new Eppendorf tubes before being stored at −80 °C. Titers of lentiviral vectors were determined by qPCR Lentivirus Titration (Titer) Kit (Applied Biological Materials, Richmond, BC, Canada, catalog #: LV900).

### Confocal fluorescence microscopy

Cells were grown on 6-well glass-bottom plates (Cellvis, catalog #: P06-1.5H-N) in complete growth medium. Cells were imaged in a precision-controlled environmental chamber using an Olympus IX83 confocal fluorescence microscope (Tokyo, Japan) under 20x or 40x magnifications. The images were captured using a galvanometer scanner. Filter settings were as follows: 359 nm (excitation) / 461 nm (emission) for DAPI, 488 nm (excitation) / 510 nm (emission) for GFP, and 579 nm (excitation) / 603 nm (emission) for mKate. Data collection and processing were performed using the software package Slidebook 5.0. All images within a given experimental set were collected with the same exposure times and underwent identical processing.

### RNA sequencing (RNA-seq)

For RNA-seq, total RNAs were harvested using the RNeasy Mini kit (Qiagen, catalog #: 74104), and sample purities were evaluated using OD260/OD280 (1.8–2.2). Genewiz Standard RNA-Seq service was employed, which requires 2 µg of total RNA in 10 µL nuclease-free water. The assays were performed on an Illumina HiSeq platform (2 × 150 bp configuration, single index), and the outputs contained ∼20 million reads per sample. All samples had mean quality scores larger than 30. For data analysis, the human reference genome (GRCh38, https://genome-idx.s3.amazonaws.com/hisat/grch38_tran.tar.gz) was used, and a pipeline consisting of HISAT2, StringTie, and DESeq2 was employed to identify differentially expressed genes (filtering conditions: adjusted p-values < 0.01, and |log2(fold-change)| > 1). For fusion protein detection, a pipeline of STAR and Arriba was used (filtering conditions: counts of split_reads1 and split_read2 > 3).

### Real-time RT-PCR

For real-time RT-PCR assays, total RNAs were extracted using the RNeasy Mini Kit (Qiagen, catalog #: 74104). First-strand cDNAs were synthesized using QuantiTect Reverse Transcription kit (Qiagen, catalog #: 205311). Next, quantitative PCR was performed using the KAPA SYBR FAST universal qPCR Kit (Kapa Biosystems; catalog #: KK4601), with GAPDH as the internal control. The forward primer (P17) for human GAPDH was 5’-AATCCCATCACCATCTTCCA-3’, and the reverse primer (P18) for human GAPDH was 5’-TGGACTCCACGACGTACTCA-3’. The forward primer (P19) for human IFIT1 was 5’-GCCTTGCTGAAGTGTGGAGGAA-3’, and the reverse primer (P20) for human IFIT1 was 5’-ATCCAGGCGATAGGCAGAGATC-3’. The forward primer (P21) for human IFIT2 was 5’-GGAGCAGATTCTGAGGCTTTGC-3’, and the reverse primer (P22) for human IFIT2 was 5’-GGATGAGGCTTCCAGACTCCAA-3’. Quantitative analysis was performed using the 2−ΔΔCt method. Fold-change values were reported as means with standard deviations (SDs).

### ChIP-seq (Chromatin Immunoprecipitation-seq)

ChIP-seq was performed as per the manufacturer’s protocol (CST, catalog #: 9003) with some modifications. Briefly, Lf-C2 and Lf-4-2 cells were treated with either DMSO or 300 nM Mithramycin A for 24 hours. Cells were crosslinked with Cross-link Gold (Diagenode, catalog #: C01019027), followed by 1% formaldehyde. Fixed cells were enzymatically digested and sonicated. Six µg of fragmented chromatin was incubated with EGR1 (Bethyl, catalog #: A303-390A), Flag (Sigma, catalog #: F1804), or IgG (CST, catalog #: 2729) overnight at 4°C. The antibody-bound chromatin was immunoprecipitated by protein G magnetic beads and was eluted. After the reverse crosslinking, the DNA was purified, and sequencing was performed at the OncoGenomics Shared Resources (OGSR) of the Sylvester Comprehensive Cancer Center. In brief, pooled libraries were sequenced using paired-end 150-base-pair chemistry on an Illumina NovaSeq X Plus instrument. The targeted average sequencing depth for ChIP-seq broad peaks was 30 million clusters per library. ChIP-seq libraries were prepared using the KAPA HyperPrep Kit (Roche, catalog #: 07962363001). Paired-end FASTQ reads were trimmed for adapters and quality using Trim Galore (v0.6.10, --2colour 20). The alignment to the human genome (hg38), peak calling, normalized bigwig, and heatmaps were performed as described^23^. Peak annotation was done with HOMER (v4.11)^24^. Differential binding analysis was performed with DiffBind (v3.12.0)^25^, using FDR ≤ 0.05 and |fold change| ≥ 1.5. A consensus peak set was generated for the C2 DMSO FLAG condition using DiffBind, and differentially bound peaks from other conditions overlapping this set were excluded (bedtools intersect -v).

### Immunofluorescence assay

SFT-739 cells were seeded at a density of 2 × 10^4^ on round glass coverslips in 24-well plates. After 48 hours of incubation, coverslips were fixed with 4% formaldehyde (Merck, catalog #: 1.00496), followed by treatment with 0.2 M glycine (PanReac AppliCHem, catalog #: A1067) to reduce background staining. Cells were then permeabilized using 0.2% Triton X-100 (Merck, catalog #: X100) in 2X PBS. Blocking was performed with 1% BSA (PanReac AppliCHem, catalog #: A2244) in 2X PBS for 60 minutes at RT. The primary antibodies for BRD4 (Abcam, catalog #: ab128874) or NAB2 (Proteintech, catalog #: 19601-1-AP) were then applied at a 1:1,000 dilution and incubated overnight at 4°C. Notably, the NAB2 antibody immunogen sequence is amino acid residues 1-320, which are contained in both the wild-type NAB2 and NAB2ex4-STAT6ex2 fusion proteins. Subsequently, Goat anti-Rabbit IgG (H+L) Secondary Antibody, Alexa Fluor 488 or 546 (Invitrogen, catalog #: A11008 and A11035), were applied at a dilution of 1:1,000 for 1 hour at RT. Nucleic acid stain DAPI (Invitrogen, catalog #: 62248) was then added for 5 minutes at 1 µg/mL. To minimize cross-reactivity between two rabbit primary antibodies, sequential staining was performed. The first antibody was applied and detected with the corresponding secondary antibody, followed by extensive washing and re-blocking. This was then repeated with the second primary antibody and its corresponding secondary antibody. After thorough washing, coverslips were mounted onto microscope slides using Mowiol mounting media. Images were acquired using a Zeiss LSM900 confocal microscope.

