## Supplemental Materials for "NAB2-STAT6 Fusion Proteins Drive Nuclear Condensate Formation and Transcriptional Reprogramming in Solitary Fibrous Tumors"

**General cloning protocols**

The Q5 High-Fidelity 2X Master Mix (New England Biolabs, catalog number: M0492) was used for all polymerase chain reactions (PCR) according to the manufacturer’s protocol. All oligonucleotides were ordered from Sigma-Aldrich and were listed in **Table S6**. The plasmids were constructed
using PCR amplification, restriction enzyme digestion (all restriction enzymes were ordered from New
England Biolabs), and ligation with T4 DNA ligase (New England Biolabs, catalog number: M0202). Gel purification and PCR purification were performed with QIAquick Gel Extraction kit (Qiagen, catalog number: 28707) and PCR Purification kit (Qiagen, catalog number: 28104). Transformations were performed using NEB 5-alpha electrocompetent *Escherichia Coli* cells (New England Biolabs, catalog number: C2987). The minipreps were performed using QIAprep Spin Miniprep kit (Qiagen, catalog number: 27106). The final plasmids were confirmed by restriction enzyme digestions and direct Sanger sequencing.

**DNA constructs**

**EF1-FLAG-mKate-NAB2ex2:** The NAB2 transcript was first PCR amplified from human HCT116 genomic DNA using primers P1 and P2, and the PCR product was subsequently cloned into the Lentiviral donor vector (lenti, unpublished results) using SpeI and MluI sites. The resulting plasmid was named Step 1. Next, the 3’-UTR of the STAT6 sequence was PCR amplified using P3 and P4 primers and subsequently cloned into the Step 1 plasmid using the BlpI and MluI sites. The resulting plasmid was named Step 2. Next, fragment 1 was PCR amplified using PCMV-mKate (unpublished results) as the template and primers P5 and P6. Fragment 2 was PCR amplified using Step 1 as the template and primers P7 and P8. Next, overlapping PCR was performed using fragments 1 and 2 and primers P5 and P8, and the resulting product was subsequently cloned into the Step 2 plasmid using the EcoRI and SbfI sites.

**EF1-FLAG-mKate-NAB2ex4-STAT6ex2:** The NAB2 fragment was PCR amplified from human HCT116 genomic DNA using primers P9 and P10. The STAT6 fragment was PCR amplified from human HCT116 genomic DNA using primers P11 and P12. Next, overlapping PCR was performed using fragments NAB2 and STAT6 and primers P9 and P12, and subsequently cloned into the EF1-FLAG-mKate-NAB2ex2 plasmid using SbfI and MluI sites.

**EF1-GFP-BRD4S:** The BRD4S sequence was PCR amplified from human HCT116 genomic DNA using primers P13 and P14 and subsequently cloned into the Lentiviral donor vector (lenti-GFP-v1, unpublished results) using BsrGI and MluI sites.

**EF1-EGR1-GFP:** The EGR1 sequence was PCR amplified from human HCT116 genomic DNA using primers P15 and P16 and subsequently cloned into the Lentiviral donor vector (lenti-GFP-v2, unpublished results) using EcoRI and SpeI sites.

The sequences were saved as EF1-FLAG-mKate-NAB2ex2.gb, EF1-FLAG-mKate-NAB2ex4-STAT6ex2.gb, EF1-GFP-BRD4S.gb, and EF1-EGR1-GFP.gb in the **Supplemental_Sequences** folder, respectively.

**Figure S1**

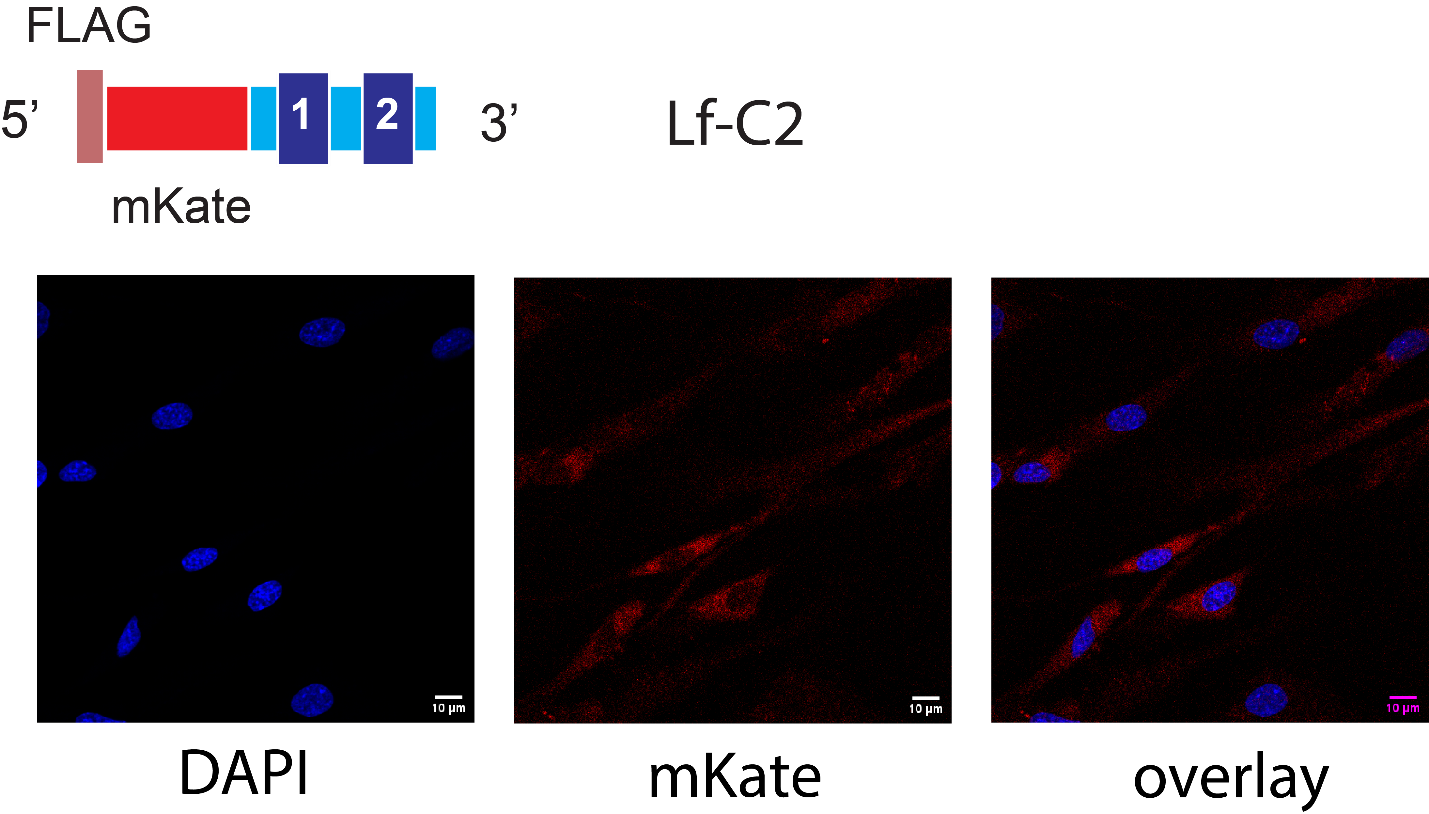

**Figure S1. Confocal fluorescence microscopy (20X magnification) showed that the control NAB2ex2 protein was mainly localized in the cytosol. (red) mKate, (cyan) DAPI**

**Figure S2**

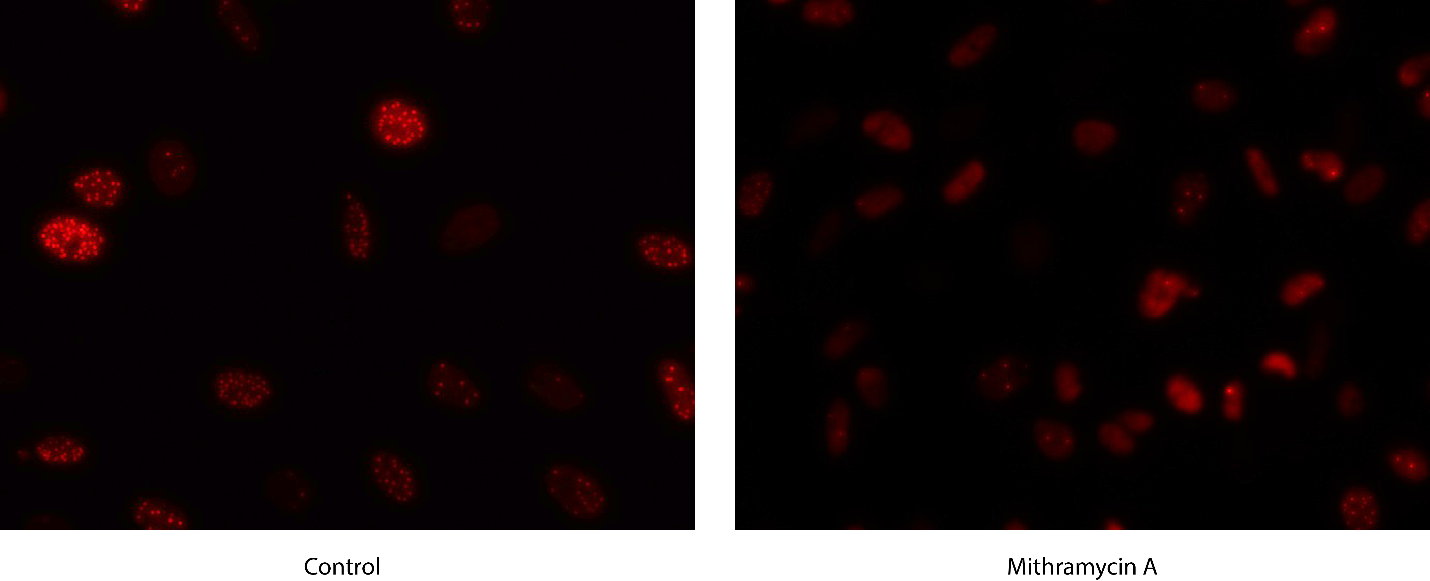

**Figure S2. Mithramycin A dissolved NAB2ex4-STAT6ex2 condensates in immortalized human lung fibroblasts.** 300 nM Mithramycin A (24 hours) effectively dissolved NAB2ex4-STAT6ex2 condensates (20X magnification).

**Figure S3**

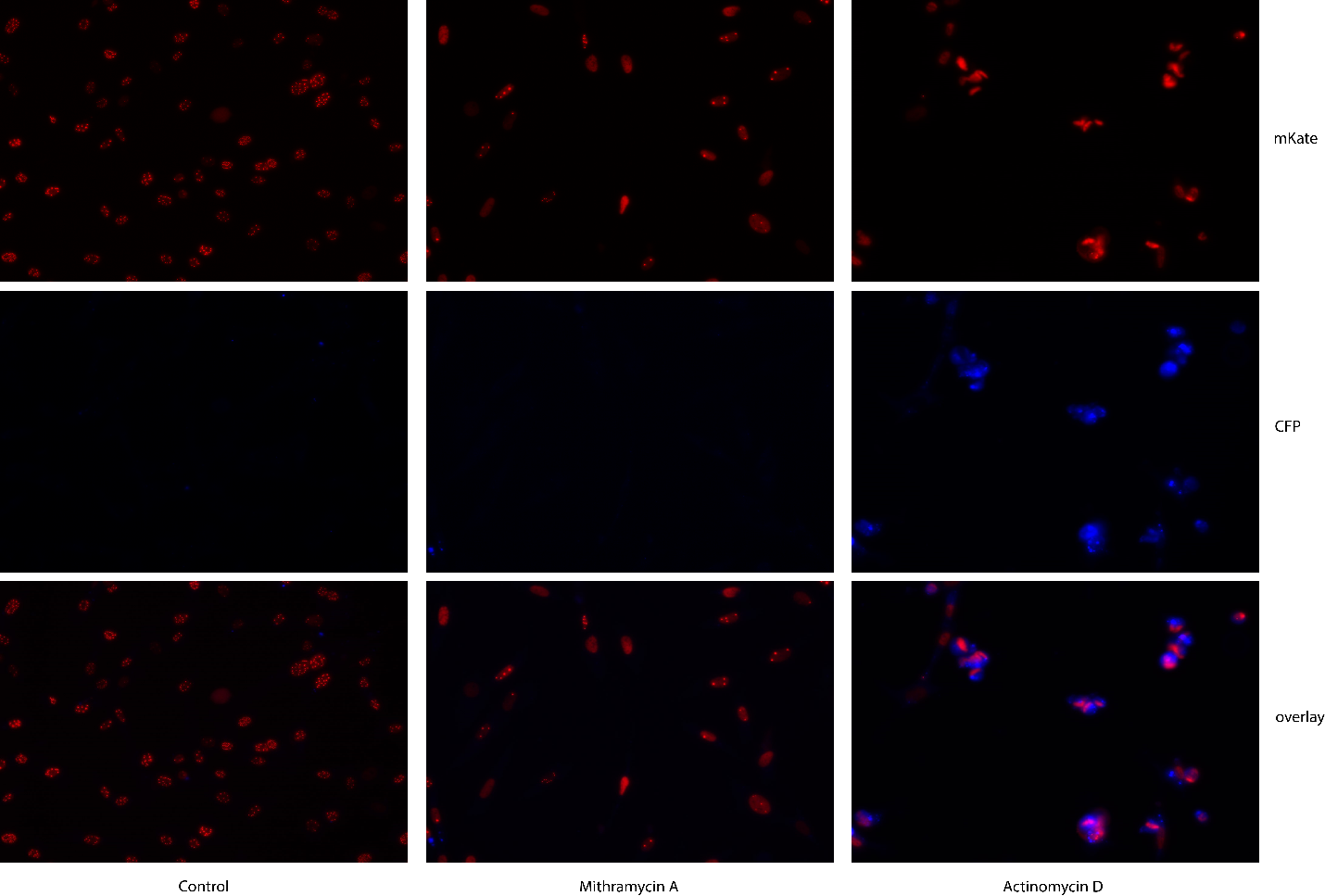

**Figure S3. Mithramycin A dissolved NAB2ex4-STAT6ex2 condensates before inducing cell apoptosis. (left)** Minimal condensate dissolution or cell apoptosis (Sigma-Aldrich, catalog number: SCT104, BioTracker NucView 405 Blue Caspase-3 dye, 5 μM) was observed in cells treated with DMSO. **(middle)** 300 nM Mithramycin A (24 hours) effectively dissolved NAB2ex4-STAT6ex2 condensates while inducing minimal cell apoptosis. **(right)** Treatment with actinomycin D (5 μΜ, 24 hours) induced condensate dissolution and cell apoptosis.

**Figure S4**

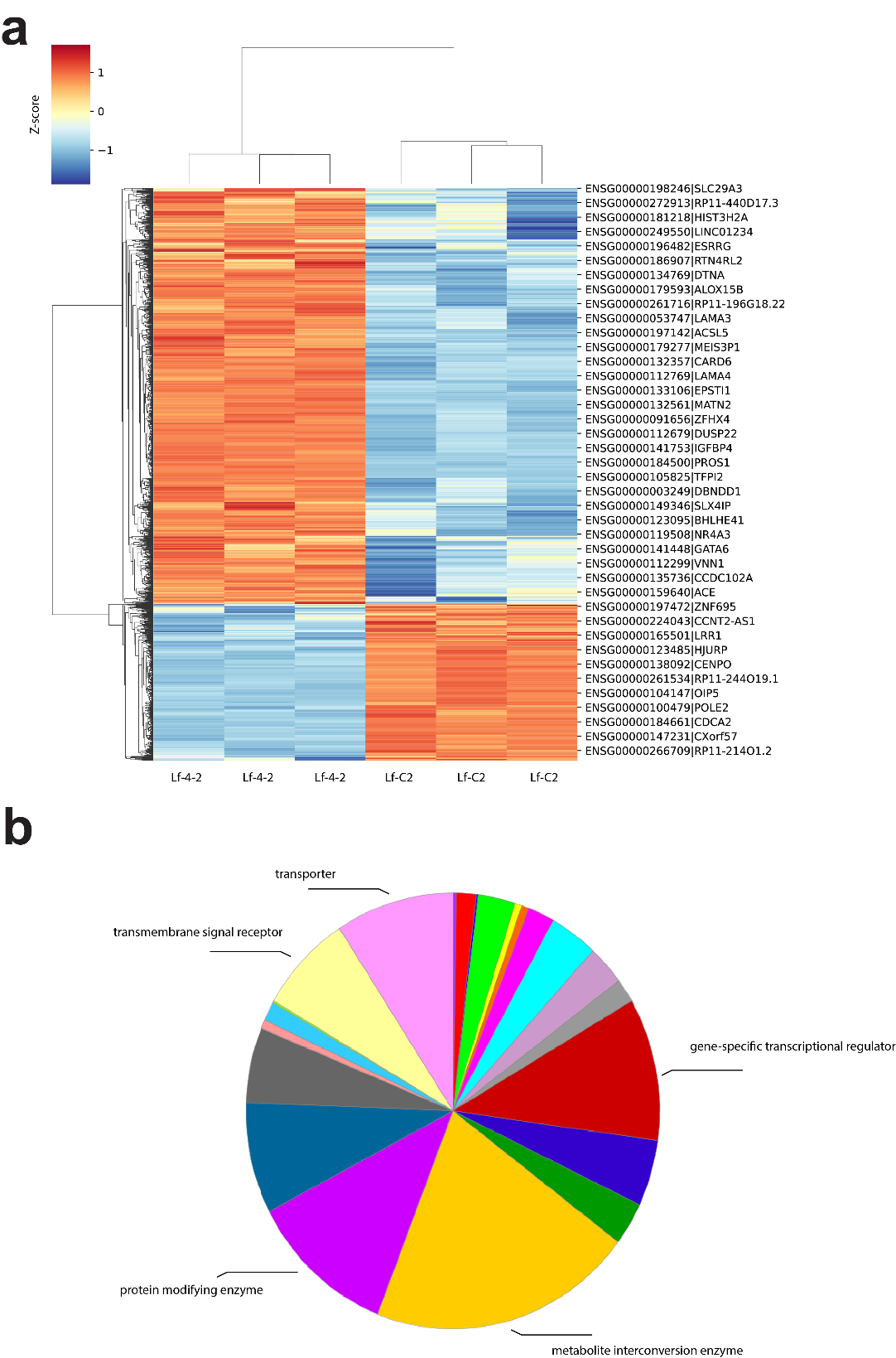

**Figure S4. RNA-seq analysis of Lf-4-2 vs. Lf-C2 cells. (a)** Heatmap showing differentially expressed genes between Lf-4-2 and Lf-C2 cells. **(b)** PANTHER analysis revealed that multiple protein classes, such as gene-specific transcriptional regulators, were enriched in candidate NAB2ex4-STAT6ex2-target genes.

**Figure S5**

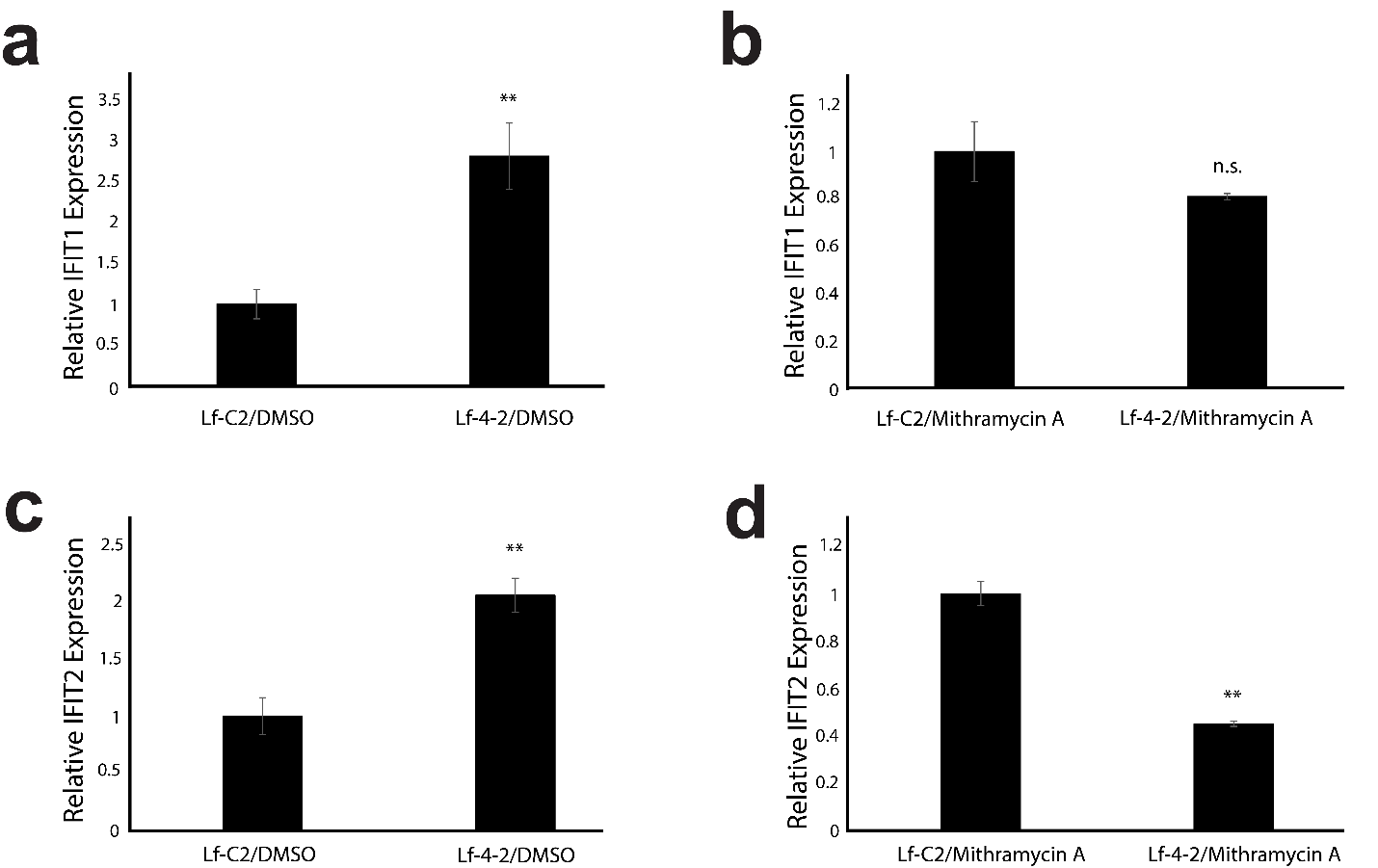

**Figure S5. Real-time RT-PCR analysis of Lf-4-2 vs. Lf-C2 cells. (a)** The expression level of IFIT1 increased in Lf-4-2 cells compared to Lf-C2 cells (2.82-fold). **(b)** The treatment of Mithramycin A reversed the overexpression of IFIT1 transcript in Lf-4-2 cells compared to Lf-C2 cells (0.81-fold). **(c)** The expression level of IFIT2 increased in Lf-4-2 cells compared to Lf-C2 cells (2.06-fold). **(d)** The treatment of Mithramycin A reversed the overexpression of IFIT2 transcript in Lf-4-2 cells compared to Lf-C2 cells (0.45-fold). For statistical analysis, two-tailed t-tests were conducted. ** p<0.01; n.s., no significant difference.

**Figure S6**

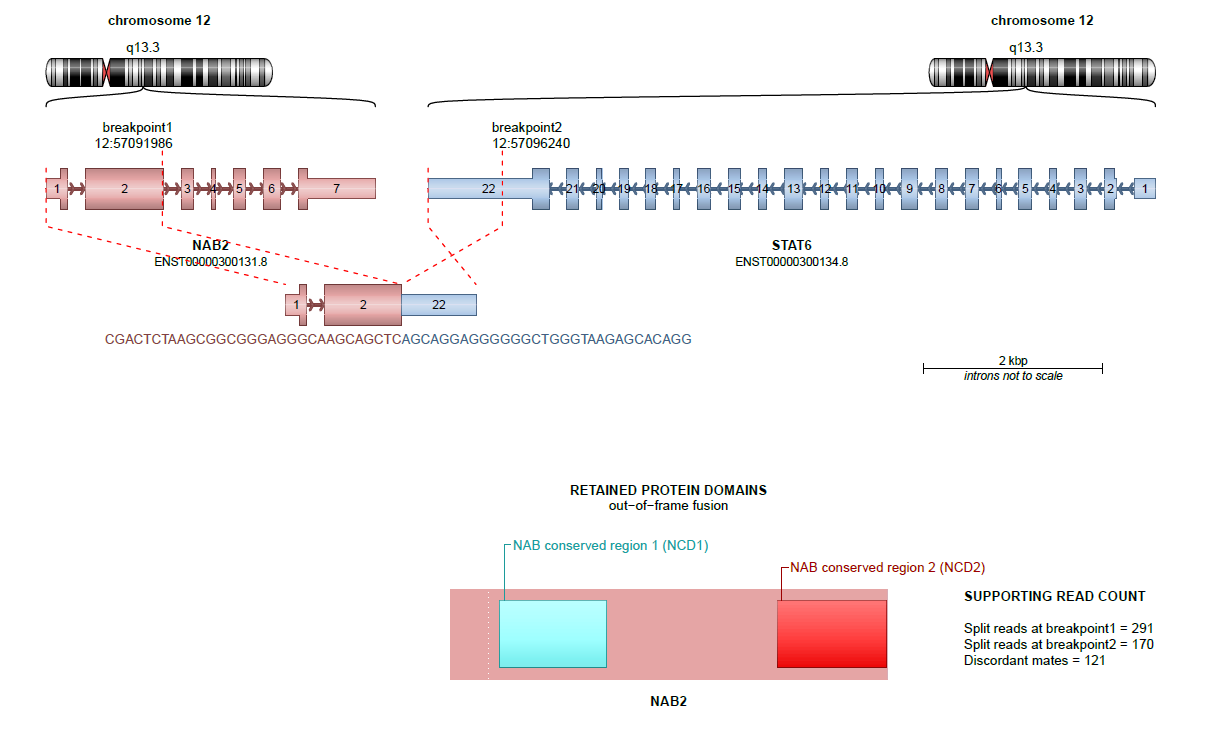

**Figure S6. RNA-seq and STAR/Arriba analysis showed that NAB2ex2-STAT6ex22 was the only fusion transcript in the genome of Lf-C2 cells.**

**Figure S7**

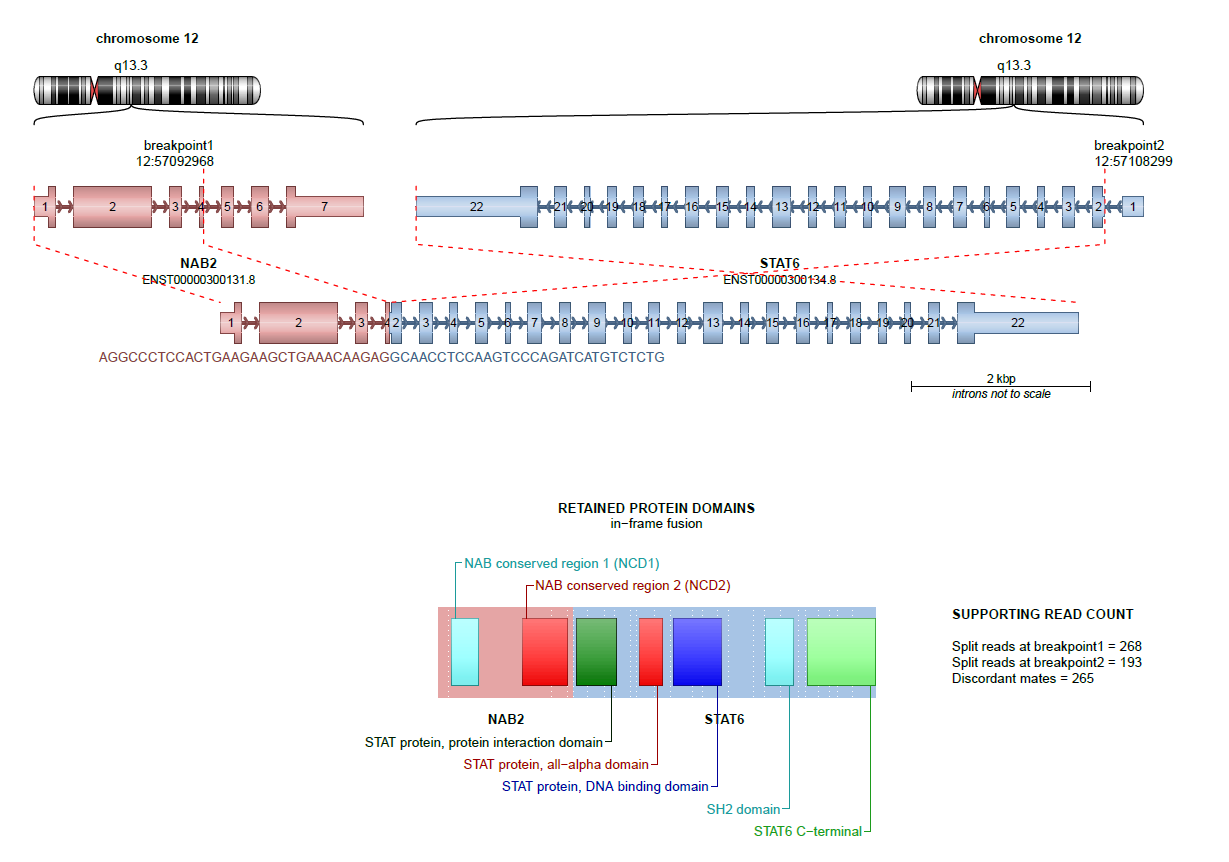

**Figure S7. RNA-seq and STAR/Arriba analysis showed that NAB2ex4-STAT6ex2 was the only fusion transcript in the genome of Lf-4-2 cells.**

**Table S1: Mithramycin A (300 nM, 24 hours) effectively dissolved NAB2ex4-STAT6ex2 condensates.**

**Table S2: Differentially expressed genes between Lf-4-2 and Lf-C2 cells from RNA-seq.**

**Table S3: Mithramycin A abolished the upregulation of expression of 856 genes that were upregulated in Lf-4-2 cells compared to Lf-C2 cells.**

**Table S4: Mithramycin A abolished the downregulation of expression of 397 genes that were downregulated in Lf-4-2 cells compared to Lf-C2 cells.**

**Table S5: The NAB2ex4-STAT6ex2 fusion transcript induced 12 EGR1-target peaks in ChIP-seq assays.**

**Table S6: The NAB2ex4-STAT6ex2 fusion transcript eliminated 20 EGR1-target peaks in ChIP-seq assays.**

**Table S7: Mithramycin A preferentially abolished the upregulation of the expression of gene-specific transcriptional regulators**

**Table S8: Primers used in this study.**
